# *De novo* designed bifunctional proteins for targeted protein degradation

**DOI:** 10.64898/2025.12.22.695915

**Authors:** Bram Mylemans, Boguslawa Korona, Amanda M. Acevedo-Jake, Ailsa MacRae, Thomas A. Edwards, Danny T. Huang, Andrew J. Wilson, Laura S. Itzhaki, Derek N. Woolfson

## Abstract

Targeted protein degradation (TPD) is a therapeutic strategy to remove disease-causing proteins by routing them to the ubiquitin-proteasome, autophagy, or lysosme machineries. For instance, proteolysis-targeting chimeras (PROTACs) are synthetic hetero-bifunctional small molecules that simultaneously bind the target and an E3 ubiquitin ligase to drive ubiquitination and degradation by the proteasome. Despite considerable success, designing such molecules is challenging and the number of currently addressable ubiquitin E3 ligases is limited. Here we demonstrate hetero-bifunctional *de novo* designed proteins as alternatives for TPD to access more targets and ligases. First, we develop a stable and highly adaptable helix-turn-helix scaffold for presenting different binding sites. Next, we use computational protein design to incorporate and embellish hot-spot-binding sites to target BCL-x_L_, plus short linear motifs (SLiMs) for KLHL20 ligase recruitment. The resulting mono- and bi-functionalised proteins bind the targets *in vitro*, and the latter degrade BCL-x_L_ in cells leading to apoptosis.

## MAIN

Over the past two decades, targeted protein degradation (TPD) has developed into a promising therapeutic strategy^1,2^. While multiple degradation machineries are under investigation^3,4^, the proteasome remains the most extensively studied^1,2,5^. For instance, molecular glue degraders^6^ and proteolysis-targeting chimeras (PROTACs)^7^ simultaneously bind a protein of interest (POI) and an E3 ubiquitin ligase (E3), inducing ubiquitination of the POI and its subsequent proteasomal degradation. Unlike classical inhibitors, PROTACs can work by engaging any region of the POI and are not limited to recognition of functional sites. Through this encounter-driven mechanism, the process can be catalytic, with multiple copies of a POI being eliminated by each PROTAC molecule^1,2,8^. As a result, lower doses of cytotoxic compounds can be used^9^. Despite encouraging progress in clinical trials^5^, bottlenecks remain for small-molecule PROTACs. The degradation efficiency is driven by ternary complex formation, which depends on multiple factors, including individual binding constants of the two recognition elements, linker lengths between them, the locations of binding sites on the targets and proximity of suitable lysine residues to which ubiquitin can be conjugated^8^. Secondly, despite the vast potential E3 repertoire, most PROTACs harness just two E3s – Cereblon (CRBN) or von Hippel-Lindau (VHL)^5,10,11^. Whilst this is changing with new E3 ligands being described^12–14^,exploring new modes of targeting and binding the POI and ligases will help advance and generalise the technology^1^. For instance, peptide-based PROTACs^15^ have the potential to overcome these limitations by using natural short linear motifs (SLiMs) as a common protein-protein interaction (PPI) modality to access more targets and E3s^7,16–18^. However, SLiMs often have weak binding affinities, making it difficult to direct precise and stable complex formation. Recent advances in protein-structure prediction and protein design might help overcome these limitations. Deep learning-based methods have accelerated the fields dramatically^19–22^, with AlphaFold2^23^ and related methods^24^ for rapid and accurate modelling, and tools such as RFdiffusion^25^ and ProteinMPNN^26^ for *de novo* protein backbone and sequence design, respectively. Computational design pipelines, either alone or in combination with experimental screening of many variants, are achieving protein binders approaching pM binding affinities^27–29^. In addition, tools like BindCraft^29^ are democratizing binder design, and allow targeting of specific regions on the POI’s surface. These successes have encouraged the field to move beyond binding a single protein. For example, proteins have been designed to inhibit multiple different POIs,^30^ or to display multiple viral epitopes^31^. Designed proteins have been fused together via a flexible linker to drive internalisation of membrane proteins^32^ or to cause degradation or stabilisation of intracellular targets^33^. Nonetheless, these strategies do not leverage the full potential of protein design, which, in principle, allows for the precise positioning of multiple binding sites to drive specific ternary complex formation.

Here, we describe a *de novo* designed protein-based PROTAC. For the POI, we target BCL-x_L_, a member of the therapeutically important B-cell lymphoma 2 (BCL-2) family. BCL-x_L_ is an anti-apoptotic protein that can be inhibited by binding helical peptide effectors leading to apoptosis^34,35^. As this interaction is well characterised with high-resolution X-ray crystal structures^36^, together with homologues such as MCL-1 it has served as a testing ground for developing methods leading to the design of many peptide-, protein- and small-molecule-based binders^37–40^. Moreover, due to interest in BCL-x_L_ as an anti-apoptotic cancer target, small-molecule inhibitors and PROTACs have been developed for it^41^; *e.g.*, a PROTAC synthetic molecule, DT2216, is currently undergoing clinical trials^42^. Like many PROTACs, this uses a VHL-binding ligand to induce ubiquitination. Recently, we have identified other E3s compatible with TPD of BCL-x_L_, including Kelch-like protein 20 (KLHL20) with an in-cell, GFP-based degradation assay (accompanying manuscript). X-ray crystal structures^43^ indicate that the KLHL20 binding-competent SLiM adopts a compact loop conformation. Indeed, binding affinity can be improved by constraining the SLiM as a cyclic peptide, and these have been used in the design of small-molecule/peptide hybrid PROTACs^17^.

To generate and validate a *de novo* protein-based PROTAC against BCL-x_L_, we develop the following design pipeline (Fig. 1a). First, using rationally seeded computational protein design^44^, we make a straightforward helix-loop-helix scaffold. Next, we add a binding site for BCL-x_L_ using hot-spot-grafting combined with ProteinMPNN optimisation and AlphaFold2 modelling and evaluation. To test the versatility of the scaffold and approach, we design and compare binders for BCL-x_L_ and MCL-1, exploring specificity with *in vitro* binding assays and structural studies. Subsequently, we graft on a SLiM (degron) for KLHL20 and optimise its binding affinity. These bindings sites were tested individually in the in-cell, GFP-based degradation assay for their capacity to act as targeting sequences or degrons. Finally, the best performing binding sequences were combined and assessed for their ability to degrade BCL-x_L_ and induce apoptosis in a lung-cancer cell line. Specifically, we compare single and bifunctional constructs with binding sites against BCL-x_L_ and/or KLHL20, and the small molecule in clinical trials, DT2216. Only the constructs with both modalities—*i.e.*, for target and E3 ligase binding—function as BCL-x_L_ degraders in cells.

**Figure 1:**
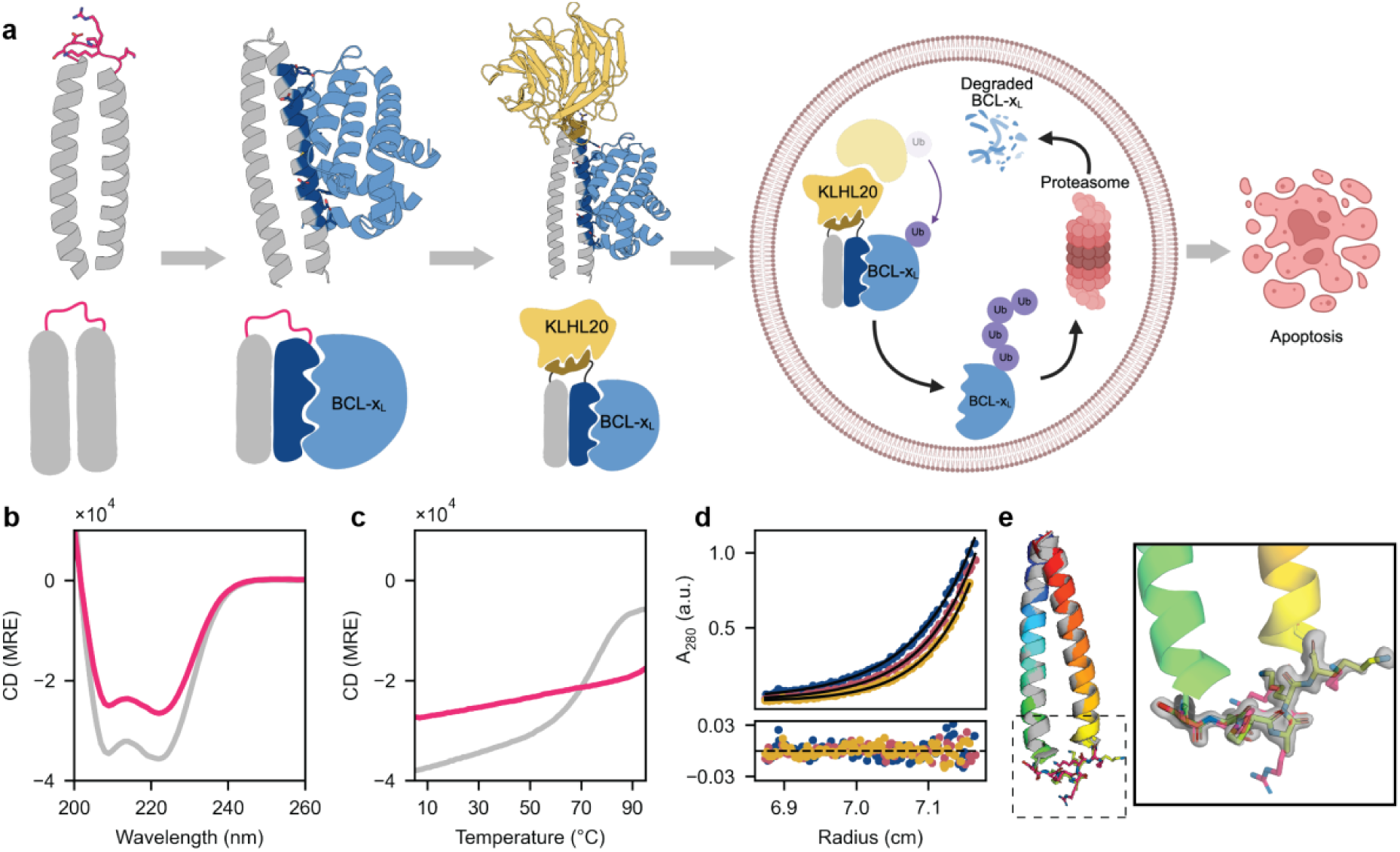
*De novo* design of a scaffold for display of multiple binding motifs. **a,** Schematic overview of this work. From left to right: helix-loop-helix scaffold (sc-apCC-2) designed based on the heterodimeric, antiparallel, coiled-coil peptides, apCC-Di-AB peptide^45^; grafting of the BCL-x_L_ binding site; addition of a second interaction site for the E3 ligase adapter, KLHL20, into the loop; and testing for BCL-x_L_ degradation and apoptosis in human cell lines. **b**, Circular dichroism (CD) spectra for apCC-Di-AB (grey) and sc-apCC-2 (magenta). CD spectra were recorded at 20 °C and 50 µM total peptide/protein concentrations. **c,** Thermal response of CD signals at 222 nm for apCC-Di-AB (grey) and sc-apCC-2 (magenta), at 50 µM and 10 µM, respectively. **d,** Sedimentation-equilibrium analytical ultracentrifugation measurements at 52,000 RPM (blue), 56,000 RPM (red) and 60,000 RPM (yellow). The ratio between experimentally fitted and theoretical mass was 0.9 over the three speeds. **e,** X-ray crystal structure of the sc-apCC-2 scaffold, electron density around the loop shown at sigma level 1 (PDB id: 9TLL; 1.5 Å). Parts of this figure were made with BioRender.

## RESULTS

### Rational design yields a highly stable display scaffold

To create a small, highly thermostable and, highly designable scaffold for displaying helical target-binding sites and additional sites for E3 recruitment, we applied our rationally seeded peptide-to-protein approach^44^. Specifically, we generated a helix-loop-helix scaffold from a previously characterised antiparallel, heterodimeric *de novo* coiled coil, apCC-Di-AB^45^. Starting with the X-ray crystal structure of apCC-Di-AB, motif searching with MASTER^46^ was used to find loops from the PDB to connect the two helices. Next, the sequence of one loop and flanking residues was optimized for the scaffold using Rosetta remodel^47^ (Supplementary Fig. 1). Initially, this designed hairpin was made by Fmoc-based solid-phase peptide synthesis. Unlike the dimeric parent peptide, this was highly thermostable by CD spectroscopy. An X-ray crystal structure at 1.5 Å resolution (PDB id: 9TLL) had an overall backbone RMSD of 0.36 Å between the design model and experimental structure, and it confirmed to the loop design. We called this new 67-residue miniprotein scaffold, sc-apCC-2, using our systematic naming system^44^.

### ProteinMPNN-embellished motif grafting gives robust binders to the POI

Previously, we have shown that dimeric coiled-coil peptides can bind to BCL-2 family proteins by grafting hot-spot residues onto the outer helical surfaces^48,49^. To improve on this, we added modern AI tools such as ProteinMPNN^26^ and AlphaFold2^23^ to the following computational design pipeline, which is depicted in Fig. 2a. To seed binder designs, first, we examined experimental structures of natural binders and targets in the BH3-only BCL-2 family and identified six hot-spot positions. To generate starting sequences for these sites, we used a small amino-acid pallet reflecting the sequences of BCL-2 effector proteins. For example, to target BCL-X_L_ binding, we started with 18 different sequence motifs (Supplementary Table 10) and grafted these onto 3 plausible locations on the solvent-exposed surface of the *de novo* scaffold. Models for each of the resulting 54 seed sequences were aligned onto 5 models for BCL-x_L_:effector complexes where the pose of effector peptide was rotated incrementally by 10° (Supplementary Fig. 2), leading to 270 models. Next, to explore the potential to extend the complementary binding surfaces, we applied ProteinMPNN-RosettaRelax protocols^26^ and varied the number of residues in the scaffold to be mutated. This yielded 609 different sequences. For each of these, AlphaFold2 was used to predict complexes with BCL-x_L_. These models were filtered and assessed based on pLDDT, different AlphaFold2 metrics, shape complementarity (Sc),^50^ and Rosetta energy (ΔG)^51^ to select candidates for experimental testing. We repeated this protocol for MCL-1 as a secondary target. While also a member of the BCL-2 family, it has a distinct selectivity profile and frequently shows increased levels as a response to BCL-x_L_ inhibition^52^.

**Figure 2:**
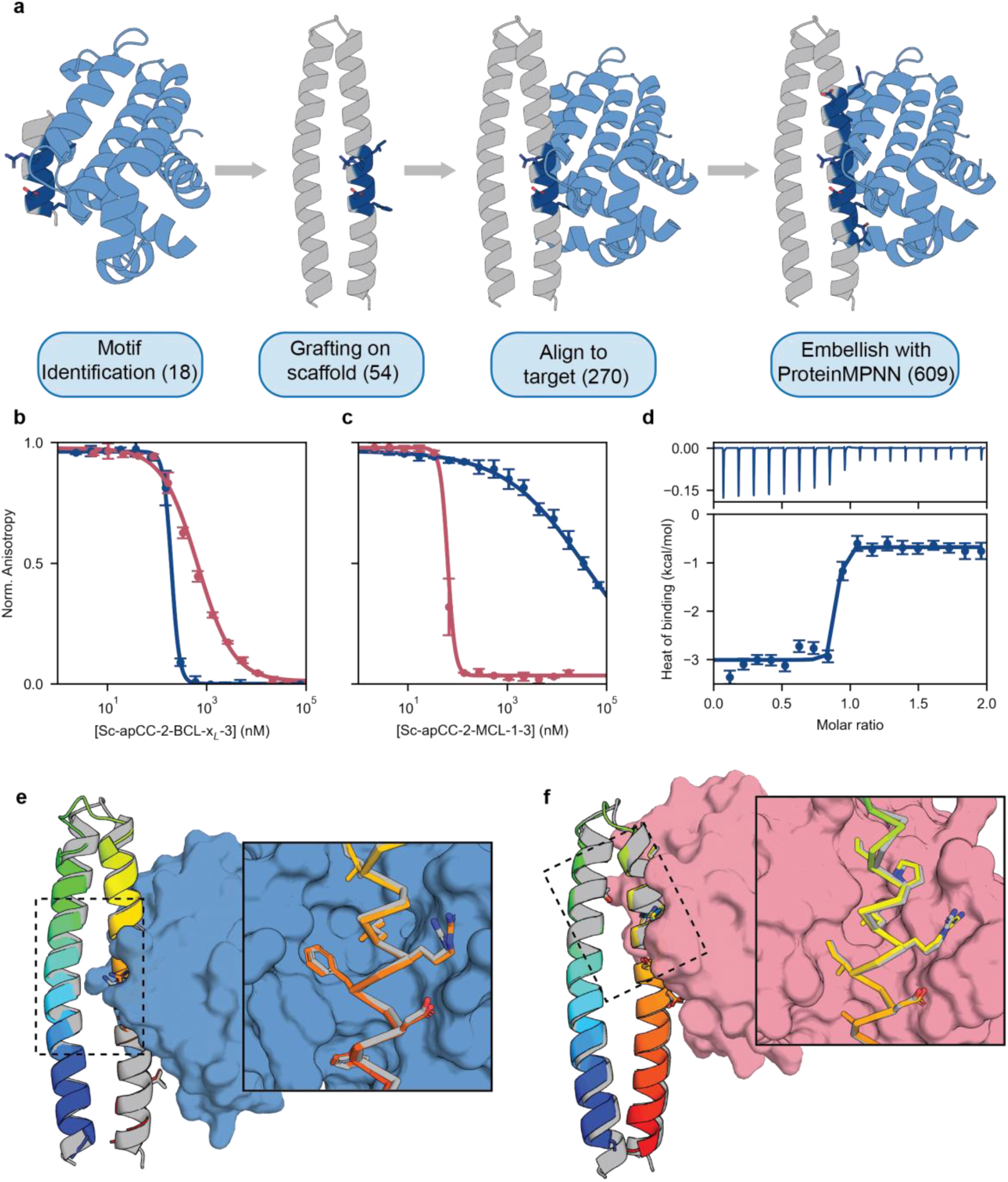
*De novo* designed binders for BCL-2-family proteins. **a**, Design schematic. From left to right: hot-spot residues selected from natural BCL-x_L_ and MCL-1 effectors were grafted onto the scaffold at all feasible surface-exposed locations of the scaffold and then aligned to the binding cleft; ProteinMPNN was used to add residues at the interface to improve binding. All final designed sequences were predicted as complexes with the target using AlphaFold2. Numbers in brackets refer to the motifs/designs at each step in the pipeline. **b**, Fluorescence anisotropy (FA) data for a BCL-x_L_ binder against BCL-x_L_ (dark blue, IC_50_: 196 ± 14 nM) and MCL-1 (red, IC_50_: 660 ± 50 nM). **c**, Similar FA data for a MCL-1 binder against MCL-1 (red, IC_50_: 64 ± 2 nM) and BCL-x_L_ (dark blue IC_50_: 40 ± 20 µM). **d,** ITC data and analysis for the BCL-x_L_ binder against BCL-x_L_ (K_D_: 3 nM. **e,** Co-crystal X-ray structure of the BCL-x_L_ binder with BCL-x_L_ (PDB id 9TLU). The experimental structure of the sc-apCC-2 component (shown in rainbow colours) is aligned with its predicted model (grey), and the experimental structure of BCL-x_L_ is rendered as a surface (blue). The zoom shows details of the interacting side chains, the non-interacting helix and side chains are not shown for clarity. **f,** Similar images to **e**, but for the co-crystal X-ray structure of the MCL-1 binder and MCL-1 (PDB id 9TLV). The scaffold components in **e** and **f** are aligned vertically to highlight the shift in the designed binding sites.

We chose 14 designs against the primary target BCL-x_L_ and 4 against MCL-1 for expression from synthetic genes in *E. coli*. All but one of these could be purified as soluble protein, and the remaining 13 were highly thermostable helical proteins (Supplementary Fig. 3). X-ray crystal structures revealed that the surface mutations were accommodated by the scaffold (PDB ids: 9TLM, 9TLN, 9TLO, 9TLP, 9TLQ, 9TLR, Supplementary Fig. 4-5). Initial binding to both targets was probed using a competitive fluorescent anisotropy (FA) assay with the natural effector peptide, BID, as the labelled competitor peptide that was displaced (K_D_ = 71 nM against BCL-x_L_ and K_D_ = 21.5 nM against MCL-1, Supplementary Fig. 6 and Supplementary Table 11)^48,49^. 12 of the expressed designs had sub-µM IC_50_ values against their intended targets, near the edge of the dynamic range for this assay (Supplementary Table 12). The MCL-1-targeting sequences were highly selective, while nearly all BCL-x_L_ targeting designs bound both targets equally.

Isothermal titration calorimetry (ITC) was used to validate BCL-x_L_ binding for a selected binder, Sc-apCC-2-BCL-x_L_-3. This binder had an IC_50_ of 196 ± 14 nM by FA, and a K_D_ of 3 nM at 37 °C by ITC (Fig. 2d and Supplementary Table 14). A 3 Å resolution X-ray crystal structure for this complex revealed that side chains in the binding interface matched the predicted conformation (Fig. 2e; PDB id 9TLU, 3.0 Å). Crystal structures of two MCL-1 binders in complex with the target were also obtained (Fig. 2f and Supplementary Fig. 7; PDB ids 9TLV, 9TLW), likewise confirming the accuracy of the designs and AlphaFold2 predictions.

### Computational negative selection improves selectivity for the POI

Presumably due to the similarity between the peptide-binding interfaces on MCL-1 and BCL-x_L_, the designed BCL-x_L_ binders showed little specificity. To improve this, we added the following computational negative-selection step as follows to identify selective designs for BCL-x_L_. AlphaFold2 was used to model the complexes of all previously designs with both target proteins (Fig. 3a). These models were then scored and ranked based on the pLDDT of the motif residues, overall pAE score, and Sc. Sequences with large differences in rank for each metric and a combined metric were selected (Fig. 3b, and Supplementary Fig. 8). In total four further designs targeting BCL-x_L_ and two for MCL-1 were expressed in *E. coli*. Five of these could be purified as soluble proteins. Their biophysical characteristics were similar to the foregoing designs (Supplementary Fig. 9). From the FA assay showed that, the BCL-x_L_ binder selected by Sc, Sc-apCC-2-BCL-x_L_-17, retained a high affinity for BCL-x_L_ and, moreover, it bound MCL-1 >1000-fold more weakly (Fig. 3c). The other designs selected using the other metrics—pLDDT and pAE, and a combination of these plus Sc—either showed little specificity or had decreased affinity for both targets (Supplementary Fig. 7 and 8). Thus, ProteinMPNN-enhanced motif grafting with some filtering—in this case based on shape complementarity scores—can generate tight and selective binders. That said, while the number of designs tested here is limited, these data suggest that current confidence-based metrics do not reliably distinguish between candidates for experimental testing. Further work is needed to assess computational designs more effectively, and to make pipelines like those outlined here more predictive.

**Figure 3:**
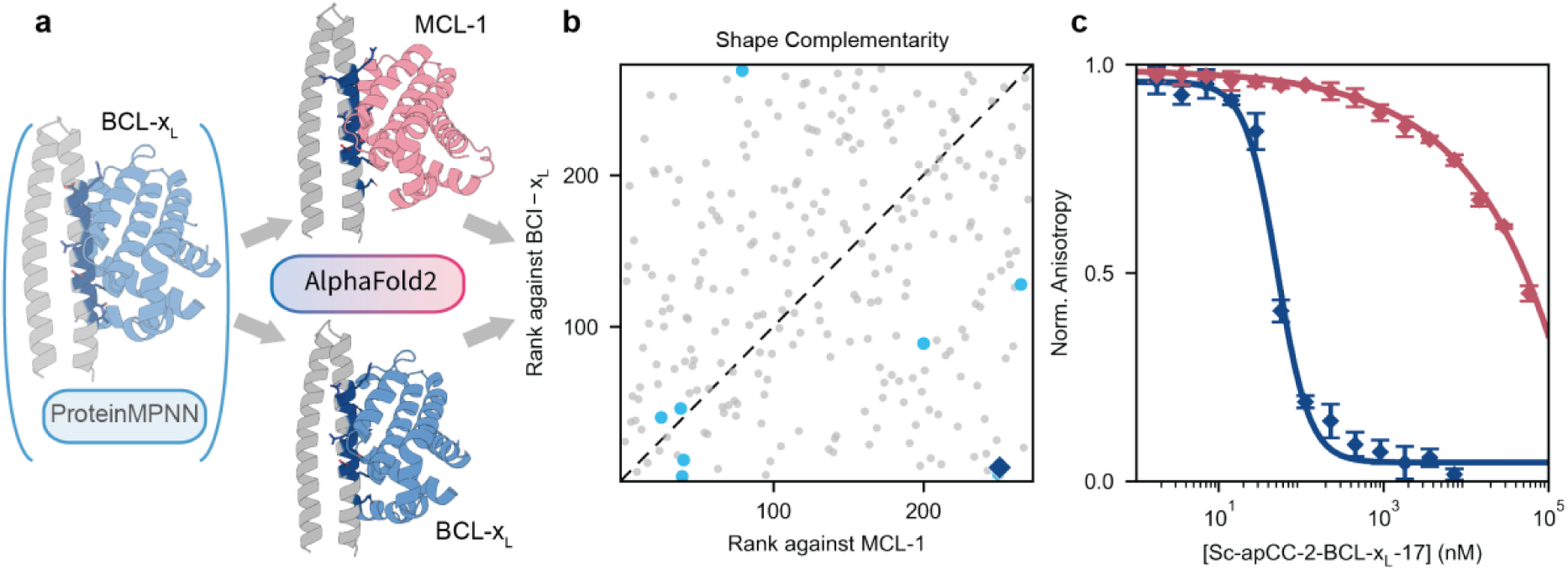
Designing selective binders for BCL-x_L_ through computational negative selection. **a,** Schematic of the selectivity procedure. After the ProteinMPNN design step against the desired target, AlphaFold2 was used to predict the complex with both targets, MCL-1 (red) and BCL-xL (blue). **b,** Designs were ranked for binding to both targets based on metrics such as shape complementarity (Sc, shown in this scatterplot; for others supplementary Fig. 6). Designs predicted to be selective for BCL-x_L_ are located in the bottom right corner. The designs selected based on Sc is shown as a dark blue diamond, and all previously tested anti-BCL-x_L_ designs as cyan circles. **c,** FA data for the BCL-x_L_ selective sequence based on Sc, binding data against BCL-x_L_ (dark blue, IC_50_: 52 ± 3 nM) and MCL-1 (red, IC_50_: 68 ± 80 µM).

### E3 targeting by grafting a SLiM for KLHL20 into the designed loop

With potent BCL-x_L_ binders in hand, we turned to E3 targeting. The SLiM for KLHL20 adopts a turn-like conformation upon binding^43^. Therefore, we hypothesised that grafting the SLiM into the loop between the two helices of sc-apCC-2 might promote an active conformation and improve binding affinity. We replaced the loop with either the natural SLiM in two different lengths (short, 5 residues; or long, 11 residues) based on previous studies of KLHL20 binding^43^, and we predicted models for the complexes with KLHL20 using AlphaFold2. Encouragingly, the positions of the SLiM hot-spot residues matched those in a known KLHL20/DAPK1 peptide complex^43^. However, the confidence of these initial predictions was low with pLDDT of the grafted motif <60. Therefore, we applied the ProteinMPNN-RosettaRelax protocol keeping the four central hot-spot residues of the SLiM fixed. This increased the pLDDT to >90 (Fig. 4a). Both the natural and optimised sequences were tested experimentally with two designs for each length. Biophysical characterisation and crystal X-ray structures confirmed that the SLiM-based loops were tolerated (PDBs id 9TLS, 9TLT, Fig. 4b and Supplementary Fig. 10 and 11). In a competitive FA-binding assay for KLHL20 binding, all designs had low micromolar IC_50_ values, (direct titration of the tracer indicated a K_D_ of 12 µM, see Supplementary Fig. 5, similar to previous reports^43^). Thus, the SLiM binding sequence can be grafted into the loop of sc-apCC-2 to yield KLHL20 binders without compromising other scaffold properties. Although the designs with the longer, 11-residue loops had higher affinities as did the ProteinMPNN-optimised designs relative to the natural sequences, the differences were small with all IC_50_ values in the low µM range (Supplementary Fig.10; Supplementary Table 13).

**Figure 4:**
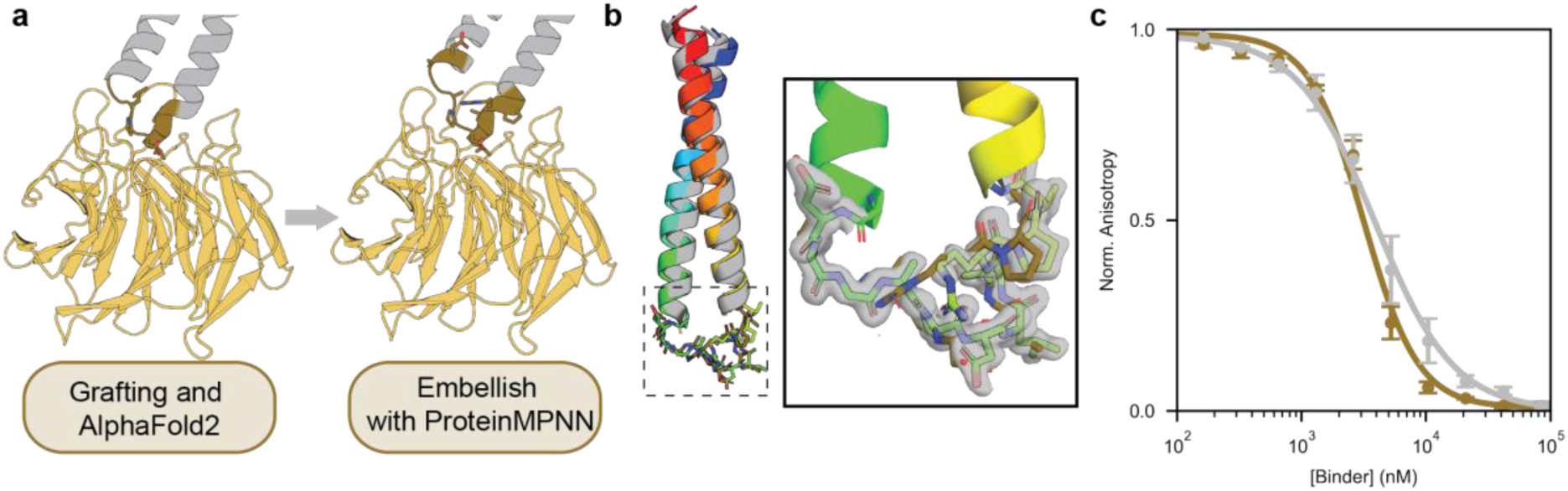
Incorporation of a SLiM for E3 ligase recruitment into the *de novo* scaffold. **a**, A natural SLiM sequence (brown) was grafted into the loop of the scaffold, and AlphaFold2 used to predict the complex with the recognition domain of KLHL20. ProteinMPNN was used to alter residues close the target. **b,** X-ray crystal structure of the sc-apCC2-KLHL20-3 (ProteinMPNN short-loop design; PDB id 9TLS), electron density shown around the loop at sigma level 1. **c,** FA data for KLHL20 binding with the natural short-loop design (grey) (IC_50_: 3.9 ± 0.2 µM) and the ProteinMPNN short-loop design (brown) (IC_50_: 3.16 ± 0.18 µM).

### BCL-x_L_ and KLHL20 binders recruit their targets in a GFP-based degradation assay

To test the capacity of the monofunctional BCL-xL and KLHL20 binders to recruit their targets, we used a previously established GFP-based degradation assay HEK293 cells (accompanying paper). The assay follows degradation of transiently expressed protein and allows a medium-throughput assessment of individual binders to inform those that could be combined in single construct. In brief, to test the BCL-x_L_ binders, HEK293 cells engineered to express a BCL-x_L_:GFP fusion protein stably were transfected with BCL-x_L_ binders fused to the full-length SPOP E3 ligase. 4 out of 8 binders showed significantly decreased GFP fluorescence compared to the negative control. To test the KLHL20 binders, the same HEK293 cells were transfected with the KLHL20 binders fused to an anti-GFP nanobody. All 4 binders lowered GFP fluorescence (see Supplementary. Fig. 12).

### Bifunctional designs are potent degraders of BCL-x_L_ and induce apoptosis

After validating their degradation capacity individually, we combined the BCL-xL and KLHL20 binding sites into a single scaffold, and we tested for endogenous BCL-x_L_ degradation in a lung-cancer cell line (A549). For this, we chose the most potent and best performing BCL-x_L_ binder from the GFP-based degradation assay, sc-apCC2-BCL-x_L_-5, and combined it with the KLHL20-binding module, sc-apCC2-KLHL20-3. These were configured in two ways (Fig. 5a): one with two binding motifs combined within a single coiled-coil module; and a second with the two monofunctional modules linked in tandem (Supplementary Fig. 13). For negative controls we used both *de novo* binding proteins individually. As a positive control we used PROTAC DT2216, which comprises small-molecule ligands for BCL-x_L_ and VHL (Supplementary Fig.14). A fluorescently labelled anti-BCL-x_L_ antibody was used to measure endogenous BCL-x_L_ levels. Neither of the separate monofunctional controls significantly decreased BCL-x_L_ levels (Fig. 5b&d. However, both bifunctional designs showed significant decreases in BCL-x_L_, just as DT2216 (Fig. 5b&d). Addition of the proteasome inhibitor carfilzomib decreased the level of degradation significantly, indicating a proteasome-depended mechanism (see Supplementary Fig. 15). Combinations of other BCL-x_L_ and KLHL20 binding sequences yield similar results, highlighting the versatility of the scaffold and the binding site design (Fig. 5d and Supplementary Fig. 16).

**Figure 5:**
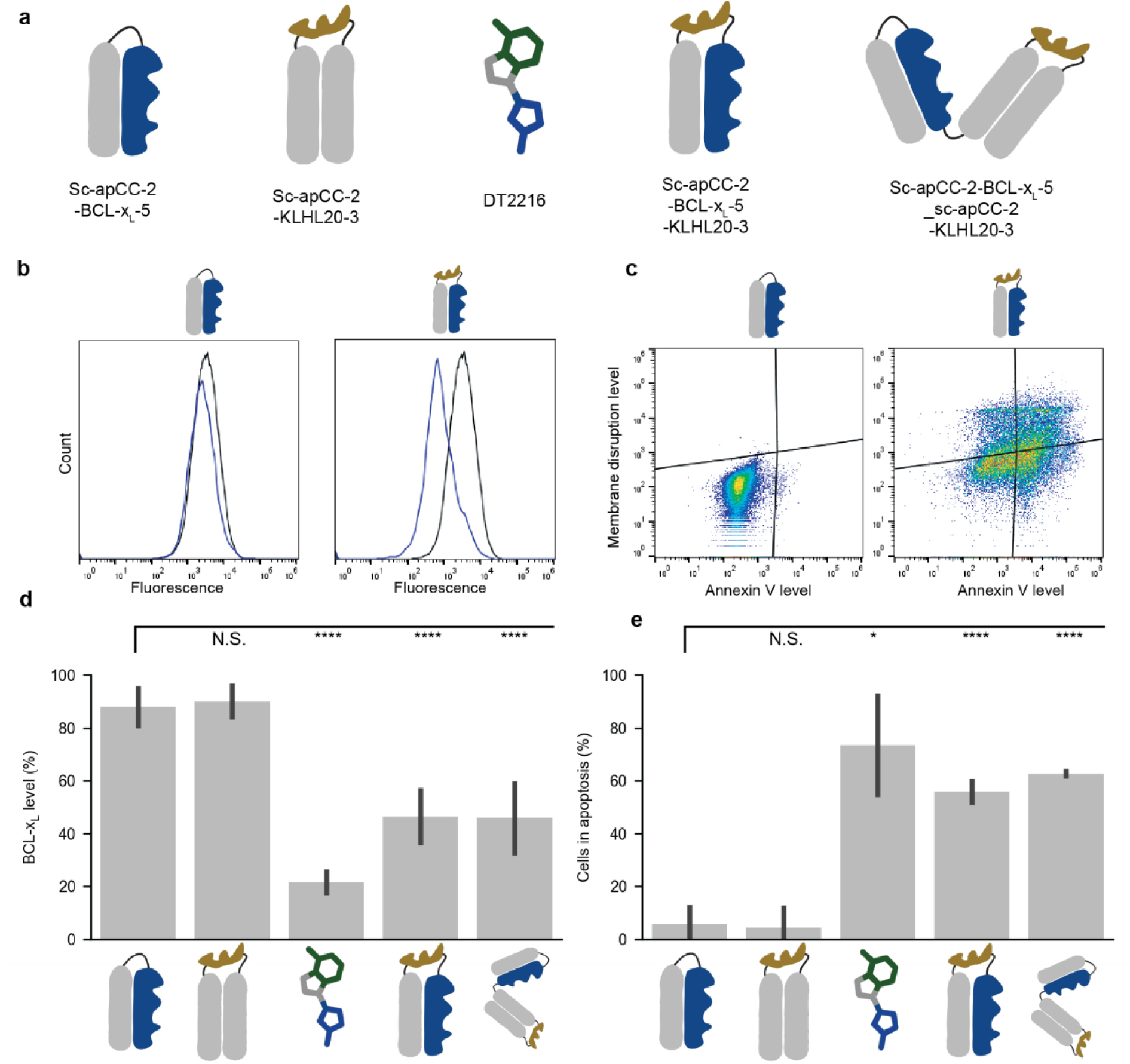
Bifunctional binders cause endogenous BCL-x_L_ degradation and apoptosis. **a,** Schematic of the molecules tested. All the designs use the same BCL-x_L_- and KLHL20-binding motifs. Negative controls are monofunctional (*i.e.*, contain only the BCL-x_L_-binding motif (blue) or the KLHL20-binding motif (brown). Small-molecule PROTAC DT2216 was used as a positive control (the chemical structure is only a cartoon representation). **b,** Flow cytometry results for the endogenous degradation, representative results for the negative control and combined binders are shown. Histograms for design (blue) and mCherry control (black). **c,** Flow cytometry results for the apoptosis assay Annexin V level on the x-axis, and DAPI signal indicating membrane disruption on the y-axis. Representative results are shown for negative control and combined binder. **d,** Degradation assay for native BCL-x_L_ in the A549 cell line. BCL-x_L_ levels were calculated by the Overton positive method, average of at least three biological repeats. Significance levels calculated with Welch t-test against the BCL-x_L_ binder. **e,** Apoptosis assay, bars show combined early and late apoptosis averaged over at least three biological repeats. Significance levels calculated with a Welch t-test against the BCL-x_L_ binder. Significance levels are represented as N.S.: P > 0.05, ∗: P ≤ 0.05, ∗∗∗∗: P ≤ 0.0001. Parts of this figure were made with BioRender.

Depleting BCL-x_L_ should increase apoptosis. To test this, we used another 2D FACS assay to measure (i) externalised phosphatidylserine indicative of early-stage apoptosis using FITC-labelled Annexin V (x-axis), and (ii) cell viability using DAPI to monitor loss of membrane integrity associated with late-stage apoptosis (y-axis)^53^ (Fig. 5c). Overall apoptosis (combined early and late) mirrored the protein-degradation assay (compare Fig. 5c&e): the individual binders did not increase apoptosis above baseline, whilst the bifunctional helix-loop-helix constructs showed significant increases, comparable to the addition of DT2216. These results were mirrored in a ApoTox-Glo™ Triplex Assay, which assesses cell viability, cytotoxicity and apoptosis events simultaneously (Supplementary Fig. 17). Combined, these experiments show that transfection of bifunctionalised scaffolds into live cells causes robust degradation and subsequent induction of apoptosis on a par with the small-molecule PROTAC.

## DISCUSSION

In summary, to develop *de novo-*protein-based PROTACs, we report the computational design of a small, highly stable protein scaffold to present multiple epitopes that engage different subcellular proteins. Using this, we introduce a binding site for the anti-apoptotic, cancer target BCL-x_L_ and another for an E3 ubiquitin ligase, KLHL20. Effectively, this produces a *de novo* protein based PROTAC that brings these proteins together in cells. Indeed, the bifunctional proteins do cause degradation of BCL-x_L_ and increase apoptosis in a cancer cell line. The designed proteins are characterised fully *in vitro* allowing them and the design methods to be evaluated.

Recently, the incorporation of multiple, independent binding sites into a single *de novo* scaffold has been reported^31^. Moreover, others have combined engineered or designed proteins to render the EndoTag and GPlad systems for clearing proteins by internalisation into cells, or via modification followed by proteolysis within cells, respectively^32,33,54^. In The PROTAC field polypeptides have long been used as modules^7^. Recently, protein design tools such as RFDiffusion^25^ and ProteinMPNN^26^ have been used to generate bifunctional unstructured peptides that bind the androgen receptor and the E3, VHL, leading to degradation of the former^55^. The design of a structured helical dimer comprising a SOS mimetic and p53 mimetic within a crosslinked dimer has been shown to engage MDM2 E3 ligase, and reduce levels of RAS^56^

Our designs are distinct from the peptide-based systems, as they are stable, folded proteins that simultaneously bind a target and an E3 ligase. Nonetheless, they still allow the flexible introduction of different binding sites by grafting onto structured helical surfaces or within the constrained loop. This promises opportunities in future to screen target—E3 pairings, and to direct the orientation of the bound proteins for tailored ubiquitination. This *de novo* design approach also offers possibilities for generating proteins that intervene in and direct other post-translational modifications such as phosphorylation^57^.

Our approach also facilitates full biophysical and structural characterisation of the designs, as demonstrated by several co-crystal structures that we report here. In turn, this helps critically assess the designs and design methods. For instance, we use motif grafting embellished with computational sequence design using ProteinMPNN followed by AlphaFold2 structure predictions. In almost all cases, the addition of residues in direct contact with the targets improves the AlphaFold2 and physical-scoring metrics. Interestingly, and as reported by others^26,27^ extending ProteinMPNN to residues outside of the binding site improves the AlphaFold2 confidence metrics but does not alter the physical scoring metrics untouched (Supplementary Fig. 19). Moreover, improvements suggested by *in silico* modelling and assessment are not necessarily reflected in experimental measurements of binding constants. For instance, for the BCL-2-family binders, extending the binding site beyond the natural, hot-spot motif proved essential to achieve better binding; however, when targeting the KLHL20-binding site there was no improvement beyond using the natural (SLiM) sequence. This raises the potential problem of false positives in design pipelines. Moreover, and aligned with recent large-scale *de novo* binder studies^58^ and other benchmarks^59^, we find that none of the AI confidence predictors or physical scoring metrics correlate well with the measured experimental binding affinities (Supplementary Fig. 20-24). Thus, in our view and corroborated by others^58–60^, the development of better *in silico* predictors of binding affinity, or simply better ranking schemes for the best binders to test experimentally are key and important challenges for the *de novo* design field.

## MATERIALS AND METHODS

### Computational loop design

The last and first six residues from the A and B chain of the previously published crystal structure of apCC-Di-AB (PDB:7Q1T) respectively were used as a search query for MASTER^46^. The original database made available with the publication was used. Results up to 0.6 Å RMSD were separated by the number of residues in between the helices. Loops of 4 residues were the most common and the lowest RMSD hit (PDB id 2XPI residues 470-473) was chosen as the starting structure for remodelling. Starting from the aligned and inserted loop, 100 Rosetta remodel^47^ trajectories were run, a total of 6 residues (the 4 loop residues and 2 flanking residues) were mutated to any amino acid but cysteine. The remaining sequence was fixed. The lowest energy model was selected for experimental testing.

### Computational binding site design

For the BCL-x_L_ binder design, a selection of 6 residue motifs based on the natural binders (BAD and BIM) were created. All motifs have the following structure Φ_1_…LΦ_2_XD.F with Φ_1_ either Y,F or I, Φ_2_ either F,I,M and X either G,S. The full list of motifs is shown in supplementary table 10. These motifs were grafted onto a modified model of sc-apCC-2, so that none of the core coiled-coil heptad positions (**a** and **d**) were altered. To accommodate the binding sites better, 4 residues were added to each helical termini, the R@**a**, D@**e** pairs were split and charges were more evenly distributed at **e** and **g** heptad positions as is the case in apCC-Di^45^. This scaffold could accommodate the motifs at 3 locations resulting in a total of 54.

These models were aligned on 5 slightly modified AlphaFold2 predictions of natural effector:protein complexes. A Rosetta script introduced a small angular shift in the effector binding angle, the resulting 5 models are shown in supplementary figure 2. Subsequently, 4 different strategies were used to modify the sequence with a ProteinMPNN-RosettaRelax pipeline: Strategy 1, no additional mutations were made; Strategy 2 non-motif, non-core and non-loop residues within 6 Å from BCL-x_L_ are allowed to mutate; Strategy 3 all non-motif, non-core and non-loop residues are allowed to mutate; Strategy 4 all non-core residues are allowed to mutate (thus including the natural motifs and designed loop). The ProteinMPNN-RosettaRelax pipeline consisted of 3 cycles of ProteinMPNN at a temperature of 0.2, grafting of the top sequence onto the helix-loop-helix and a short relax following the InterfaceRelax2019 script. After the final relax run, Rosetta scores including shape complementarity and interface ΔG were calculated. In addition, an AlphaFold2 prediction was made of each final sequence with and without a pre-generated MSA of BCL-x_L_. Designs were filtered based on pLDDT (>85 as a monomer and >90 as a complex) and a final selection was made based pLDDT, pAE, ΔG or shape complementarity. All selected sequences with their respective strategy and selection method are shown in supplementary table 4.

For MCL-1, the same method was employed but only the MPNN implementation in strategy 2 was used. Instead of strategy 1, the entire previously described binding site^48^ was grafted onto the modified sc-apCC-2 model. Selected sequences are shown in supplementary table 3.

For KLHL20, the previously designed loop was replaced with the natural SLiM sequences - AGLPDLVGA-(short) or -AGLGLPDLVAKYNGA-(long). AlphaFold2 models were predicted with a pre-genrated MSA for KLHL20. Subsequently the same three cycle ProteinMPNN-RosettaRelax pipeline was used keeping the middle-PDLV-of the SLiM fixed.

All code and inputs are available from the labs Github page.

### Computational selectivity screening

For the BCL-x_L_ and MCL-1 selectivity screening, AlphaFold2 predictions were made of the final sequences of strategy 2 with the opposite target’s MSA. In addition, the final sequences were grafted on sc-apCC-2 and target complexes and subsequently relaxed and scored by Rosetta. For pLDDT, pAE and Shape Complementarity the rank of each sequence against each target was determined, sequences with the largest difference in rank favouring the desired target were selected. All code and input are available from the labs Github page.

### Solid-phase peptide synthesis and purification

All peptides used in this study (see Supplementary Table 1) were synthesised following standard Fmoc automated microwave solid-phase peptide synthesis procedures with a Liberty Blue (CEM) as descripted before^44,49^. N,N’-diisopropylcarbodiimide (DIC) and Oxyma Pure in N,N-dimethylformamide (DMF) were used as activators and a 20% (v/v) solution of morpholine or piperidine in DMF was used in the deprotection step. Peptides with acetyl (Ac) caps were treated with pyridine or diisopropylethylamine and acetic anhydride. 5-carboxyfluorescein (FAM) coupled peptides were prepared by coupling of 6-(amino)-hexanoic acid and FAM with the standard DIC and Oxyma Pure in DMF mixture. Peptides were cleaved from the resin with a trifluoracetic acid (TFA):water:triisopropysilane mixture in 95:2.5:2.5 (v/v) ratio at room temperature. Subsequently, cleaved peptides were precipitated with ice-cold diethyl ether.

The freeze-dried peptides were dissolved in a water/acetonitrile mixture and purified with reverse-phase HPLC using Water and acetonitrile with 0.1% (v/v) TFA as solvents. Peptide purity was assessed by analytical HPLC (see Supplementary Fig. 25, 31, 33, 35, 37), and matrix-assisted laser desorption/ionization-time of flight (MALDI-TOF) mass spectrometry (see Supplementary Fig. 25, 30, 32, 34, 36).

### Protein expression and purification

The genes encoding the designed binders and target proteins (BCL-xL, MCL-1) were cloned into a pet28a vector with Kanamycin resistance. E. coli BL21 DE3 cells were transformed with the plasmids (see Supplementary Tables 2-7 for protein sequences). Cells were grown in Luria Broth at 37° C until an optical cell density of 0.6 was reached. Protein production was induced by addition of isopropyl-β-D-thiogalactoside and cultures were kept at 20 °C overnight. Afterwards, the cells were harvested by centrifugation and frozen until further use.

For purification of the binders, the cells were resuspended in binding buffer (25 mM Tris, 300 mM NaCl, 10 mM imidazole pH 8.0) and lysed by sonication. The lysate was clarified by centrifugation. The supernatant was applied on a His-trap column equilibrated with binding buffer and washed with (30 mM imidazole. The protein was eluted with 300 mM imidazole. Subsequently, the imidazole was removed by a PD-10 desalting column and incubated overnight with TEV protease at room temperature. After His-tag cleavage, the protein solution was reapplied on the His-trap column and flow-through was collected. This flow-through was applied on a HiLoad 16/600 superdex75 column equilibrated in biophysics buffer (25 mM Tris, 150 mM NaCl, pH 7.4). The peak containing the monomeric proteins was collected and concentrated for later use.

For BCL-xL and MCL-1 protocols were followed as previously described^49^. The cells were resuspended in lysis buffer (25 mM Tris, 500 mM NaCl, pH 8.0 cOmplete, Mini, EDTA free protease inhibitor cocktail, Roche and 5 mg lysozyme and 3 mg DNAse, Roche). Afterwards, they were sonicated and clarified via centrifugation. The soluble fraction was applied on a His-trap HP column (Merck) pre-equilibrated with wash buffer (10 mM imidazole, 25 mM Tris, 500 mM NaCl, pH 8.0). The column was then washed four times with increasing concentration of imidazole up to 100 mM, before a final elution with 400 mM imidazole. The protein solution was dialyzed overnight together with His-Ulp1 to remove the SUMO-tag. The next day the solution was applied on the His-trap HP column and the flow-through was collected. This flow through was collected and passed-through size exclusion chromatography with a HiLoad 26/60 Superdex 75 column. This column was previously equilibrated with 20 mM Tris 250 mM NaCl, 0.5 mM DTT, 2.5% glycerol, pH 8.0. The fractions containing the monomeric protein were pooled and concentrated before storage for later use. Protein purity and quality was assessed by mass spectrometry (see Supplementary Fig. 26-27) and CD (see Supplementary Fig. 29).

For KLHL20 expression and purification, Rosetta™ (DE3) Competent Cells (Novagen) were transformed with a KLH20 DNA construct (pNIC28-Bsa4-6his-KLHL20(303-605). Cells were grown to OD600 of 0.6 at 37 °C, before the temperature was lowered to 18 °C and protein expression was induced with 0.4 mM IPTG overnight. The cells were pelleted by centrifugation and resuspended in lysis buffer (50 mM HEPES pH 7.5, 500 mM NaCl 5% glycerol, 0.5 mM TCEP, 5 mg lysozyme, 3 mg DNAse and 1 cOmplete, Mini, EDTA free protease inhibitor cocktail tablet). Afterwards, the cells were sonicated and clarified by centrifugation. The supernatant was applied to 5 mL HisPur Ni-NTA resin (Thermo Fisher), preequilibrated with binding buffer (50 mM HEPES pH 7.5, 500 mM NaCl, 5% glycerol, 0.5 mM TCEP) and left rotating at 4 °C overnight. The flow through was collected using a peristaltic pump and the beads were washed four times with increasing imidazole concentration (10 mM −100 mM). Subsequently, the protein was eluted with 300 mM imidazole in the same buffer. The eluted protein was dialyzed overnight with TEV protease, before being applied again on the Ni-NTA beads. The cleaved KLHL20(303-605) was eluted with the same elution buffer. The protein was dialyzed into 50 mM HEPES pH 7.5, 300 mM NaCl, 0.5 mM TCEP, 2.5% (v/v) glycerol and concentrated before storage for later use. Protein purity and quality was assessed by mass spectrometry (see Supplementary Fig. 28) and CD (see Supplementary Fig. 29).

### Circular dichroism spectroscopy

Circular dichroism (CD) spectroscopy data were collected on a JASCO spectropolarimeter instrument fitted with a Peltier temperature controller. The binders were prepared to 20 µM concentration in 20 mM Tris 150 mM NaCl, pH 7.4 buffer unless otherwise stated in figure legends. Data were collected in a 1-mm quartz cuvette with a 200 nm to 260 nm wavelength. The following instrument settings were used: band width, 1 nm; data pitch, 1 nm; scanning speed, 100 nm/min; response time 1 s. Each spectrum is the background subtracted average of 8 scans. For the thermal denaturation scans, the circular dichroism signal was followed at 222 nm wavelength while increasing the temperature from 5 °C to 95 °C at a ramp rate of 1 °C/min. BCL-x_L_, MCL-1 and KLHL20 were measured in 10 mM K_2_HPO_4_ adjusted with NaOH to pH 7.4 at 50 µM protein concentration. The spectra were normalized to mean residue ellipticities (deg cm^2^ dmol^-1^res^-1^) by normalizing for concentration of peptide bonds and the cell path length using equation 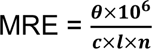 with *θ* the measured difference in absorbed circularly polarized light in mdeg, *c* the concentration in µM, *l* the path length in mm and *n* the number of amide bonds in the proteins.

### Analytical size exclusion chromatography

Analytical size exclusion chromatography was performed on a JASCO HPLC system with a superdex 75 increase 10/300 column. 20 µL of protein solution at a concentration of 100 µM in 20 mM Tris, 150 mM NaCl, pH 7.4 was injected at 0.5 mL/min. The UV absorbance at 230 nm was recorded, the background was subtracted and the normalized signal was plotted in function of elution volume.

### Analytical ultracentrifugation

Sedimentation equilibrium experiments were performed on a Beckman Optima X-LI analytical ultracentrifuge with an An-60-Ti rotor (Beckman-Coulter) and data was fitted using SEDPHAT software^61^ to single-species model. Monte Carlo analysis was performed to yield 95% confidence limits. A peptide concentration of 100 µM in PBS (8.2 mM sodium phosphate dibasic, 1.8 mM potassium phosphate monobasic, 137 mM NaCl, 2.4 mM KCl, pH 7.4) was used. The sample was spun at 52000, 56000 and 60000 RPM after 8-hour equilibration at each speed, the measurement at each speed was duplicated.

### Competitive fluorescence anisotropy assay

The competitive fluorescence anisotropy experiment were performed following previously described procedures^49^. Each experiment was run in triplicate and the fluorescence anisotropy measured using a CLARIOstar plus plate reader (BMG Labtech), with excitation at 482nm (bandwidth = 16nM), emission at 530nm (bandwidth = 40nm) and polarised dichroic mirror at 504nm. For the direct titration assay, stock concentrations of 1mg/ml peptide were made in DMSO and then further diluted to a working stock of 100nM in FA buffer (20mM tris, 150mM NaCl pH7.4 1mM DTT 0.01% triton x100) for use in the assay. For MCL-1 and BCL-x_L_ experiments, the target protein concentration was fixed at 150 nM and tracer concentration fixed at 25 nM. For KLHL20 experiments, KLHL20 concentration was fixed at 8 µM and 83.3 nM tracer concentration. The competitor protein was titrated in. Perpendicular (P) and parallel (S) intensities of the control wells were averaged and deducted from each corresponding samples to give the corrected values, P_corr_ (P_corr_ = P_sample_ – P_av. control_) and S_corr_ (S_corr_ = S_sample_ – S_av. control_). The total sample intensities (I) and anisotropies (r) were calculated with the follow equations: I = (2 x P_corr_) + S_corr_, r = (S_corr_ – P_corr_) / I

The points for each competition curve were normalised between 0 and 1. The obtained values were plotted against competitor concentrations and fit to a logistic function of the form Y = A2 + (A1 – A2) / (1 + (x/IC_50_)^p^) to determine the half-maximal inhibitory concentration (IC_50_) of the peptide competitor (where A2 is the upper asymptote and A1 the lower asymptote).

### Isothermal Titration Calorimetry

Isothermal titration calorimetry (ITC) experiments were carried out using a MicroCal PEAQ-ITC system (Malvern Panalytical) at 37 °C. A stock solution of BCL-xL was dialyzed into ITC buffer overnight (20 mM Tris, 150 mM NaCl pH 7.4), and the same buffer was employed to dissolve a solid portion of the tested binder. For each experiment, initially, the cell was filled with 275 µl of binder (20 μM) and the syringe was loaded with the BCL-xL at 200 µM concentration. An initial injection of 0.4 μL was then followed by 18 injections of 2 μL, every 150s, with a constant syringe rotation speed of 750 rpm throughout. The signal for the enthalpy of dilution of the protein into buffer was independently measured in a control titration experiment and then subtracted from the experimental binding traces. NITPIC^62^ was used to integrate the injections. The data was then fitted with SEDPHAT software^61^ to extract K_D_, ΔG°, ΔH° and -TΔS° using an adjusted single-site binding model.

### Crystallisation trials

Protein crystals were grown using a sitting-drop, vapor diffusion method with commercial sparse matrix screens. 0.3 µL of a concentrated protein solution or 1:1 mixture of proteins in case of the binders was mixed with 0.3 µL of the screening solution in MRC 96 well plates. The plates were incubated at 20 °C until protein crystals were observed. The crystallisation condition for each structure can be found in Supplementary table 15. The crystals were mounted and transferred to a cryoprotecting solution made of the screen supplemented to 25% glycerol before flash freezing in liquid nitrogen.

The proteins were diffracted at Diamond light source or the European Synchrotron Radiation Facility. Data processing was done on the beamline with either XDS^63^ or Dials^64^. Aimless^65^ was used for scaling. Molecular replacement was done with PHASER^66^ using AlphaFold2 models. Phenix Refine^67^ was used for refinement of the structures using translation/liberation/screw (TLS) parameters for each chain. Data and refinement quality statistics can be found in supplementary table 16.

### Cell culture

All cell culture procedures were carried out according to standard procedures. In brief, all cell lines were maintained at 37 °C in an atmosphere of 5% CO₂ and ≥95% humidity. HEK293T cells were cultured in Dulbecco’s Modified Eagle Medium (DMEM; Gibco™) supplemented with 10% foetal bovine serum (FBS), while A549 cells were grown in Nutrient Mixture F-12 Ham medium (Merck) supplemented with 10% FBS. DNA transfections were performed using the cationic lipid reagent TransIT-2020 (Mirus Bio).

Stable GFP–Bcl-x_L_ cell lines (see Supplementary Table 8) were generated in a previous study (accompanying paper) through antibiotic selection followed by sorting of live GFP-positive cells for subsequent propagation.

### Flow cytometry assay of degradation of GFP-BCL-x_L_

BCL-x_L_ binders were synthetized in pcDNA3.1Hygro(+) plasmid (GenScript) in such a way that they are preceded by a single FLAG tag and followed by the SPOP E3 ligase and mCherry with a T2A self-cleaving peptide sequence. The KLHL20 binder constructs were in the same plasmid preceded by a FLAG tag and anti-GFP nanobody and followed by the same mCherry T2A combination. Full protein sequences are found in Supporting Table 8.

The flow cytometry assay was performed as previously descripted (accompanying paper). Briefly, on day 1, cells were seeded at a density of 300,000 per well supplemented DMEM without antibiotics. On day 2, cells were transfected following the manufacturer’s instructions, with a ratio of 0.5 µg DNA to 1.5 µL transfection TransIT-2020. On day 4 (48 hours after transfection), cells were washed with PBS, trypsinized, resuspended in FACS buffer and stained with DAPI (1 µg/mL) prior to flow cytometry on a Attune NxT instrument (ThermoFisher Scientific). The data was analysed with FlowJo v10. Events were gated based on fluorescence of cells stably expressing GFP-BCL-x_L_ and transfected with an mCherry-only control (quadrant Q2). Degradation of BCL-x_L_ was indicated by reduced GFP fluorescence intensity and a shift to Q1. Results represent at least three independent experiments and are expressed as the percentage of remaining GFP-BCL-x_L_ (100 x Q2/(Q1 + Q2) ± standard deviation). Statistics were calculated using a two-sided Welsch’s T-test. Significance levels are represented as N.S.: P > 0.05, ∗: P ≤ 0.05, ∗∗: P ≤ 0.01, ∗∗∗: P ≤ 0.001, ∗∗∗∗: P ≤ 0.0001.

### Flow cytometry assay of endogenous BCL-x_L_ degradation

The protein constructs for the endogenous degradation assays were in the same pcDNA3.1Hygro(+) plasmid (GenScript). Constructs had N-terminal FLAG tag and were followed by mCherry preceded by a T2A self-cleaving peptide.

The flow cytometry assay was performed as previously descripted (accompanying paper). A549 cells were cultured as above and fixed in 4% formaldehyde 48 hours after transfection or 48 hours after treatment with 1 μM concentration of DT2216 (#37311, Cayman Chemical). Then, following a PBS wash the cells were permeabilized with cold methanol for 15 minutes on ice and incubated overnight at 4 °C with BCL-x_L_ (54H6) Rabbit mAb (Alexa Fluor 488 Conjugate) antibody and a Rabbit IgG Isotype control, Alexa Fluor 488 Conjugated (Cell Signaling, BS-0295P-A488-BSS-100uL Startech Scientific Ltd, respectively). The cells were washed and resuspended in PBS for flow cytometry measurements. For the proteasome inhibitor experiments cells were treated with 20 μM carfilzomib for 24 hours prior to measurements (Carfilzomib (PR-171), #A1933, APExBIO). The data was analysed with FlowJo v10. Results were plotted as events of BCL-x_L_ Alexa 488 channel. The percentage of cells that lost fluorescence was calculated by subtracting out the mCherry control using the Overton positive method. Data are from at least 3 independent experiments Statistics were calculated using a two-sided Welsch’s T-test. Significance levels are represented as N.S.: P > 0.05, ∗: P ≤ 0.05, ∗∗: P ≤ 0.01, ∗∗∗: P ≤ 0.001, ∗∗∗∗: P ≤ 0.0001.

### Flow cytometry apoptosis assay

The same DNA constructs as in the endogenous degradation assay were used. The flow cytometry apoptosis assay was performed as descripted above (accompanying paper). Briefly, 48 hours before flow-cytometry measurements A549 cells were transfected with degraders or treated with DT2216 at 1 μM (#37311, Cayman Chemical). Then, cells were detached, washed with PBS and incubated for 30 minutes to allow recovery. Afterwards, the cells were stained with FITC-conjugated Annexin V (BioLegend) in accordance with the manufacture’s protocol except for a longer 45-minute incubation time. Next, the cells were washed in 1x binding buffer and resuspend in 300 µL of binding buffer containing DAPI (BD Pharmingen). Cells, undergoing early apoptosis have externalized phosphatidylserine binding to Annexin in a calcium-dependent manner. Cells in late-stage apoptosis lose membrane integrity and get stained by DAPI. Apoptosis was calculated as the sum of cells in early and late stages and expressed as percentage of total cell count.

### ApoTox-Glo™ Triplex Assay

Viability, cytotoxicity and apoptosis assays were performed using the Promega kit according to the provided protocols, with the exception of additional filters to eliminate the mCherry fluorescence background; fluorescence filter wavelengths were 485 ± 10 nm for excitation and 520±20 nm for emission for the cytotoxicity measurements, and 400 ± 20 nm for excitation and 505±20 nm for emission for the viability measurements. Luminescence readings at 545±40 nm were used for the apoptosis measurements.

## ACKNOWLEDGMENTS

This work was primarily supported by grants from the BBSRC (BB/V006231/1, BB/V006703/1 BB/V008412/1, and BB/V008412/2), with additional support from BB/V003577/1, BB/V003577/2 and BB/Y007816/1. This project, BM, and DNW have been supported by the Novo Nordisk Foundation through the NNF Center for Protein Design at the University of Copenhagen (NNF25SA0105927). DTH was supported by Cancer Research UK core funding (A29252). X-ray diffraction experiments were supported by Diamond beam time awarded under proposal number MX34438, MX31440 and MX37593 and ESRF beamtime awarded under proposal number MX2373. We thank Alex Bullock for providing the plasmids encoding the KLHL20 plasmid. We thank Simon Coulton and Andrew Lovering for assistance with crystallographic data acquisition. We thank Diana Gimenez for preparation of the KLHL20 tracer used in FA KLHL20 competition assays. Some Figure elements were created with BioRender (https://BioRender.com/8n9fh89, https://BioRender.com/eg0oktl, https://BioRender.com/l2psuaa).

